# Diel foliar moisture content recovery time occurs soon after midnight in *Eucalyptus species* of koala (*Phascolarctos cinereus*) habitats

**DOI:** 10.64898/2025.12.10.693569

**Authors:** Ivan Kotzur, Ben D. Moore, Matthias Boer, Marta Yebra, Kara Youngentob

## Abstract

Many herbivores are reliant on the foliar moisture content (FMC, the water mass as a percentage of the dry matter mass), of feed tree leaves, which may vary through the diel cycle. This includes koalas (*Phascolarctos cinereus*), who have heightened reliance on FMC during high temperatures. However, it is unknown how, and to what degree, FMC varies through the 24-hour period. In this experiment we measured live leaf FMC every three hours over the diel period by calibrating a near-infrared spectroscopy (NIRS) model from leaf-surface reflectance, for tree species associated with koala habitat. Coinciding with reflectance measurements, trees in a glasshouse were exposed to normal and extreme climatic conditions including heatwaves and a drought, while collecting concurrent climate variables and soil moisture content. Our analysis of FMC timeseries’ of glasshouse trees found the diel range of leaf FMC was 23 % dry weight (DW, i.e. 19 % fresh weight) on average, between a minimum and maximum diel range of 10 %DW and 46 %DW, as predicted by an accurate spectral regression model (r^2^=0.96, RMSE=9.29 %DW). Overall, FMC minima occurred at 13:30 and FMC had recovered to initial levels (7:30) by 1:30 the following morning, though near-recovery levels were reached earlier by 19:30. This result shows that during a single day and night koala tree species may vary the proportion of leaf moisture by about one fifth, on average, and the timing of canopy moisture recovery may discourage foraging in the evening on temporarily dry species, during hot conditions.

## 1. Introduction

Koalas (*Phascolarctos cinereus*) are arboreal marsupial folivores that are largely dependent on pre-formed dietary water from the foliage they consume. They also produce metabolic water during the metabolism of dietary macronutrients and adipose tissue, and these two sources are sufficient to meet requirements during most conditions (Ellis et al. 2010, Degabriele & Dawson 1979). During hot conditions, koalas require an increased water intake to support the elevated rates of evapotranspiration via sweating and panting which is used to lose heat and maintain thermal balance. This is particularly so when temperatures increase above ∼30 ^O^C (Degabriele & Dawson 1979). If foliar water content is inadequate during these conditions, koalas may have to adjust behaviour to reduce their need for water or increase their water intake (Briscoe et al. 2014, Mella et al. 2019). This may be done by foraging for moist leaves, drinking free-standing water and/or increasing food intake to increase the overall volume of water intake. The latter strategy can be limited by the toxicity of koala forage, the size of their stomach and the maximum passage rate of digesta (Beale et al. 2018). Furthermore, eating incurs a heat penalty because of the thermal increment of feeding. While animals have some capacity to absorb environmental heat during the day until cooler temperatures prevail (Turner 2020), evapotranspiration demands generally fluctuate according to air temperature and relative humidity (Adam et al. 2020), with peaks at midday or in the afternoon.

Given the dependence of koala hydration on foliar moisture content (FMC), the timing of FMC changes relative to the timing of feeding and temperature changes may be critical for thermoregulation. FMC is the leaf water content on a dry weight basis:

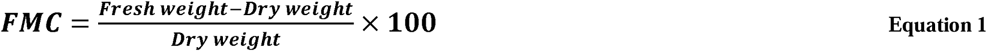

For instance, leaf FMC of 120% is 1.2 times as much moisture as dry matter. Koalas are largely but not completely nocturnal and feeding tends to occur as separate meals in between other activity (Nagy & Martin 1985). Most feeding occurs during the evening, night and early morning (Marsh et al. 2013). Temperature maxima occur during the day, and during heatwaves elevated temperatures persist into the night (Adam et al. 2020). Koalas must dissipate heat during the day, and during the night if excess body heat has been maintained. In the latter case, ability to use panting and sweating may be constrained by water intake depending on the timing of variation in forage FMC. This is particularly so if nights of elevated minimum temperatures occur in succession, limiting the effectiveness of passive cooling each night.

Koala feed trees are mostly of the genus *Eucalyptu*s, with species preferences varying regionally and within populations due largely to nutrition and toxicity (OEH 2018, Moore et al. 2014). Wide variation in FMC can occur amongst preferred species, values as high as 222 %DW have been recorded in forests of *Eucalyptus cambageana* in tropical north-eastern Australia (i.e. more than twice as much water than dry matter, Ellis et al. 2002). In contrast, FMC can be as low as 75 %DW in semi-arid floodplain woodlands of *Eucalyptus brownii* (Munks et al. 1996). Some studies have considered seasonal differences in the same trees, with preferred forage changing by 5, 12 or 18 %DW (Hume & Esson 1993, Melzer 1994, Ellis et al. 1995). These results may be increased or decreased by inclusion of the differences between young, fleshy and old, sclerified leaves. Regardless, the diel variation in FMC has rarely been documented.

*Eucalyptus* species differ in drought tolerance and in the extent to which different physiological processes are deployed to maintain water balance. Foliar moisture content is a function of leaf structure, which controls the maximum FMC value, soil moisture available to the tree, and osmotic and elastic changes in the leaf that control water content and water potential relations (Nolan et al. 2020). Therefore, research effort tends to focus on water potential, this is part of the reason for a lack of diel leaf water data. The fluctuations in leaf structure, i.e. dry matter content occur at large, seasonal timescales (Griebel et al. 2023), not relevant to the diel period, however there maybe diel changes caused by carbohydrate cycling from the leaf (Gessler et al. 2007). Other common physiological measures of plant water content include relative water content, the ratio of water content to saturated water content, however full turgidity rarely occurs within habitats and the measure is not comparable to other nutritional components of forage made on a dry mass basis. Environmentally, FMC is controlled by the atmospheric demand for water according to the vapour pressure deficit (VPD), which increases with air temperature and with declining relative humidity in the atmosphere (Nolan et al. 2022). Leaf FMC is also regulated by soil moisture availability; this varies at a longer timescale compared to VPD which varies alongside temperature within diel cycles (Ehrenberger et al. 2012). The effect on FMC of increases in atmospheric CO_2_ concentration and associated, expected improvements in plant water use efficiency (i.e. carbon gain per amount of water lost, Soh et al. 2019), remain unknown. We expect that leaf FMC will increase at higher CO_2_ concentrations because stomatal regulation for productivity can be tighter. Many plant ecophysiology studies measure water status of leaves at two time points (e.g. water potential, Nolan et al. 2020, Bartlett et al. 2012, Zolfaghar et al. 2015), of which the maximum is recorded at midday, due to the higher atmospheric demand corresponding to daily maximum temperatures (i.e. higher VPD), and the minimum is recorded at pre-dawn when temperatures are low and relative humidity increases. Measures of FMC at finer temporal resolution will indicate if there are lags between leaf water content and leaf water potential, and will provide an absolute measure of diel variations in tree water relevant to herbivore foraging.

In fire ecology, FMC is also important as a measure the availability of forest fuels for fire, where higher availability corresponds with dryer forests (i.e. lower FMC, Nolan et al. 2016). Research using satellite remote sensing is underway into the diel trends in FMC, specifically, using optical sensors with high temporal sampling frequency to measure sub-daily changes in fuel conditions (Yebra et al. 2018). Also, efforts have begun to measure plant water content using microwave satellite data in plant physiology and fire ecology (Konings et al. 2021).

Most remote sensing estimates of FMC, or similar metrics, are usually taken at one time of day which is determined by the satellite orbit and site location. Thus, remote sensing sampling time may not coincide with the timing of parameters of interest, such as FMC maxima and minima. The Sentinel-2 satellite has been used to measure fuel conditions in fire research (Yin et al. 2021), however extreme diel FMC may not occur at ∼10:30 when the satellite passes the study area in Qinyuan County, China. If diel variation is of relevance to a study, then the interpretation of a single time-point satellite measure may confound results.

Hence, it is necessary to establish a diel timeseries from leaf-level measurements to reveal any differences between the measurement condition and the condition of interest (e.g. minimum diel FMC). Furthermore, it will provide useful calibration data that allows for the adjustment of such estimates from satellites by applying interpolations from the diel trend of leaf measurements.

Near-infrared spectroscopy (NIRS) is a common method, which is similar to hyperspectral remote sensing, and has wide applicability including in assessing the nutritive value of animal forage which does not necessarily require the destruction of samples (Foley et al. 1998). This approach uses the interaction of light, both reflectance and absorbance, with biochemical constituents to predict the relative concentrations of those constituents. To make predictions, statistical calibration models need to be developed. These are most commonly partial least squares (PLS) regressions between the Visible-Near InfraRed (Vis-NIR) reflectance spectra of samples, collected non-destructively, and reference values of the constituent of interest, which are often measured destructively (Moore et al. 2010). Accurate and robust calibration models can then be used to non-destructively predict the constituent of interest in samples, including in attached, live leaves. Few studies have used NIRS calibrations to estimate foliar moisture content, i.e. water on a dry weight basis, although others have estimated foliar moisture on a fresh weight basis or relative to maximum moisture contents (Ma et al. 2019, Yang et al. 2017). Even fewer have measured FMC of live leaves, with grinding being a common preparation of samples (Gillon et al. 2004). Such preparation excludes the opportunity for monitoring leaves *in situ* or across a timeseries.

In this experiment we aimed to monitor the magnitude and timing of diel leaf foliar moisture content of several koala food trees, while manipulating the water supply and climate to imitate a range of environmental conditions, specifically heatwaves and ecological drought. Environmental variables monitored included air temperature and relative humidity (and thence vapour pressure deficit), soil moisture and solar radiation. We aimed to understand both the effect of time on leaf FMC under varied environmental conditions and to obtain a trend that could be used to adjust instantaneous FMC measures to other diel timepoints. Initially, we aimed to estimate FMC of live leaves using NIRS to rapidly, non-destructively and intensively sample the diel trend of FMC in multiple attached, live leaves.

In this study we hypothesised that minimum leaf FMC of the eucalypts will occur during the early afternoon (H_1_), due to maximum atmospheric demand on trees for moisture. We also expected that FMC would recover to previous morning levels at the time closest before dawn (H_2_), when the atmospheric demand on trees for water has reduced and the trees and leaves have had the longest time to refill with soil water. Under drought conditions, we expected delayed leaf FMC recovery time into the night (H_3_) as the lower water supply will reduce the capacity for leaf refill.

## 2. Methods

### 2.1 Eucalyptus trees

Ten trees from each of five *Eucalyptus* species were chosen based on koala preference, climate of origin and nursery availability. The trees were germinated in 3 different nurseries around the Western Sydney region, between 7 and 12 months before the experiment, resulting in trees of heights from 1.7m to 2m at the time they were moved into the glasshouse. Each tree was in a 300mm pot (25 L). *Eucalyptus tereticornis*, *Eucalyptus moluccana* and *Eucalyptus sideroxylon* are considered favoured koala feed trees, while *Eucalyptus crebra* and *Eucalyptus fibrosa* can be used as feed or shelter trees (OEH 2018). *E. crebra* and *E. sideroxylon* usually grow farther inland, in drier environments, compared to the other species (Slee et al. 2020). In this way a range of climates of origin were introduced, which effects leaf water relations (Ngugi et al. 2003), possibly including the diel patterns. All trees were allowed to acclimatise for at least 2.5 months in the glasshouse before the experiment.

### 2.2 Glasshouse conditions

The trees were randomly assigned to each of two glasshouse bays and two benches therein. In each bay, five trees from each of five species were equally treated to increasing drought over six weeks and to two heatwaves, one prior to the drought, in well-watered conditions, and one at the end of the drought (Table 1). Temperature and relative humidity, in the controlled glasshouse (Argus, Titan) were monitored using a built-in omni-sensor (Argus, SEN-OSM/CO2), which also records values of all the variables. Trees in one glasshouse bay were treated to elevated CO_2,_ of 630 ppm, adjusted constantly, compared to 430 ppm in the ambient chamber. In the well-watered condition, watering consisted of two periods daily from an automated drip system (10 min at both 9:00 and 21:00, Holman Industries, C01605). The system fed tap water via poly pipe and two drip tubes to each tree, these tubes were placed in the same position around the tree in the soil of each pot. During the droughts the period length was reduced so that two targets of soil volumetric moisture content were progressively reached: 12-14 % then 8-12 % for the daily average of each potted tree. Soil moisture content was measured using 30 cm reflectometer probes inserted at an angle because of pot height and in the same positions in each pot relative to the drip tubes and tree (Campbell Scientific, CS616 with Campbell Scientific, CR1000 logger). Measurements of two leaves on each tree (*n* = 100) were taken during eight different diel periods (i.e. days), every three hours, beginning after sunrise and ending before dawn, resulting in four each of diurnal and nocturnal timepoints. Photosynthetically active radiation (PAR) was recorded throughout the experiment (Apogee, SQ-421X-SS). Diel measurement periods occurred during a heatwave or at varying proximity to a heatwave, including prior to droughting, to capture a range of conditions of both atmospheric demand and soil water supply.

**Table 1.** Glasshouse temperature and humidity (i.e. vapour pressure deficit [VPD]) settings for the experiment. RH indicates the relative humidity.

| Hours | Temperature (°C) |  |  | VPD (kPa) |  |  |
| --- | --- | --- | --- | --- | --- | --- |
|  | Back-ground / acclimatisation | Heatwave #1 | Heatwave #2 | Background/ acclimatisation (RH=60 %) | Heatwave days 1-4 (RH=50 %) | Heatwave day 5 (RH=30 %) |
| 00-04 | 16 | 26 | 26 | 0.72 | 1.67 | 2.34 |
| 04-06 | 16 | 26 | 26 | 0.72 | 1.67 | 2.34 |
| 06-08 | 18 | 29 | 29 | 0.82 | 1.99 | 2.78 |
| 08-10 | 21 | 33 | 33 | 0.99 | 2.50 | 3.49 |
| 10-12 | 24 | 39 | 39 | 1.19 | 3.46 | 4.85 |
| 12-14 | 28 | 43 | 43 | 1.50 | 4.28 | 5.99 |
| 14-16 | 28 | 43 | 43 | 1.50 | 4.28 | 5.99 |
| 16-18 | 24 | 39 | 39 | 1.19 | 3.46 | 4.85 |
| 18-20 | 20 | 35 | 35 | 0.93 | 2.79 | 3.90 |
| 20-00 | 17 | 29 | 29 | 0.77 | 1.99 | 2.78 |
| Average | 20 | 33 | 33 | 0.99 | 2.65 | 3.70 |

**Table 2.**
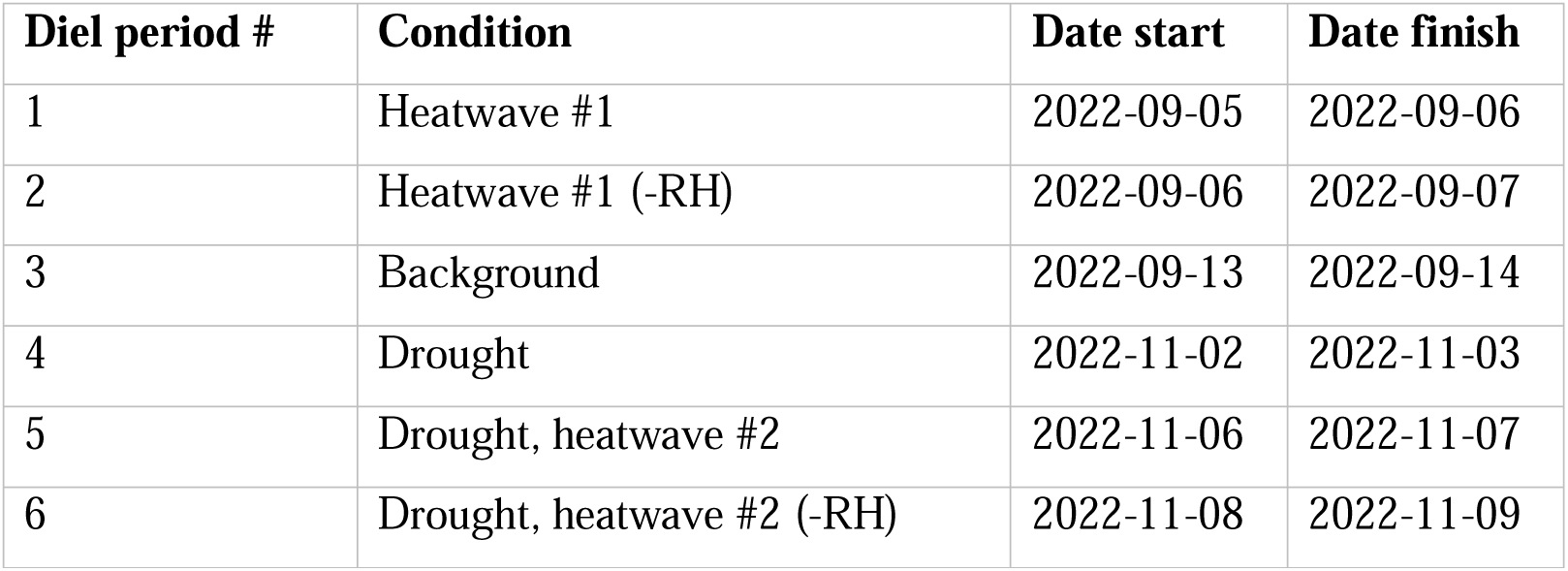
Dates of the diel periods and associated glasshouse, environmental conditions in which spectra were collected.

Manipulations included reducing humidity (thereby increasing VPD, Equation 2), using the glasshouse air conditioner, on two days during heatwaves. VPD was calculated as

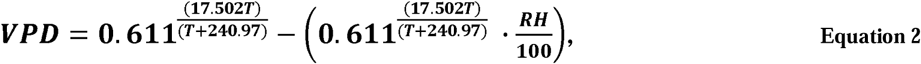

where *T* is air temperature as °C in the first term (i.e. saturation vapour pressure) and *RH* is relative humidity of the air as a percentage in the second term (i.e. actual vapour pressure), and the result is in units of kPa. Measurement periods were conducted between end of July and end of November 2022.

### 2.3 Leaf spectral measurements

Measurements of foliar moisture during the diel sampling periods were based on non-destructive scans of reflectance spectra taken using a leaf clip with a limited-damage lamp attached to a Spectral Evolution RS-3500 spectrometer. For each tree, two leaves were tagged and repeatedly measured throughout the experiment, capturing temporal rather than inter-leaf differences. All the leaves were mature and appeared healthy, however some leaves died during the experiment and a nearby leaf, similar in age and ontogeny was chosen as replacement when visible browning had occurred in the original leaf. Three spectra per leaf, from representative leaf areas (no browning, galls *etc*) within the 4mm window of the leaf clip, were collected from the adaxial surface and averaged before the leaf spectrum was used for calibration or prediction of FMC. The leaf midrib was avoided, capturing the reflectance spectrum only of leaf lamina except in instances where narrow leaves made this impossible (e.g. *E. crebra*). To collect samples that were close in time to specific diel points and because of the time required for sampling, measurements began 30 min before each timepoint and ended 30 min after that time, with all samples being assigned the central timepoint (e.g. [7:00 … 8:00] = 7:30). All times were recorded in Australian Eastern Standard Time (GMT+10), and we accounted for daylight savings time changes during the experiment in all data streams.

### 2.4 NIRS modelling of foliar moisture content

To calibrate models to predict FMC from spectra, repeated measurements of FMC and Vis-NIRS spectra were taken from excised leaves as they dried on a bench. These leaves were cut, with petiole intact, from the trees in the glasshouse during ambient conditions prior to the diel sampling periods. These leaves were from 15 trees, three of each species, and four per tree (two each of young and mature leaves). Leaves were sampled prior to dawn because leaf moisture was expected to be higher at this time and then placed into brown paper bags in a dark container with an ice pack. After transporting leaves from the glasshouse to the lab, leaf fresh weight and three spectra per leaf, were measured (0.001 g precision and same spectrometer as above). Exposing the leaves to the laboratory atmosphere, the measurements were repeated every ∼2 h, four more times, including twice the following day after being stored again in the cool container overnight. All samples in bags were then placed in a laboratory oven, set to 105°C, for 24 h. Afterwards, samples were transferred to a desiccator until the dry weight was measured. The leaf FMC was then calculated for every sampling point during the dry down following Equation 1, where *dry weight* remained constant as *Fresh weight* varied between repeat measurements. As well as these leaves, which were used as calibration and independent validation samples in partial least squares (PLS) models (see below), another set of leaves were collected at the end of the experiment to use in a second, independent validation of the models. This set was collected on one day both at pre-dawn and midday, to test the model performance at different FMC values, and were processed in the same way as the main FMC-spectra dataset.

To do the PLS modelling, the ‘plantspec’ software package in the R environment was used, which is based on the ‘pls’ package (Griffith & Anderson 2019, Mevik & Wehrens 2007). First, outliers were identified amongst the spectra of the calibration dataset. All spectra were inspected visually, per sample (i.e. three spectra), and then the Mahalanobis distance (Saptoro, Tadé and Vuthaluru 2012) was calculated across the whole set. Spectra were filtered according to squared Mahalanobis distance with a 95 % cutoff (d.f. = 2500) from a chi-square distribution. This was done using the software package ‘waves’ in the R environment (Hershberger et al. 2021). A mean of the multiple spectra per sample (n ≤ 3) was calculated and matched with the associated leaf-level FMC value. To create a subset for independent validation of the FMC model, 20 % of the input dataset was split off using the Kennard-Stone algorithm, this function maximises the Euclidean distance of the spectral wavelength vectors and so aims to make the subset representative of the whole dataset (Kennard & Stone 1969). An optimisation was undertaken to select the spectral regions and transformations that produced the best partial least squares regression models. Spectra regions tested were 400-600, 600-800, 800-1200, 1200-1600, 1600-2000 and 2000-2400 nm, and transformations tested were first and second derivatives, constant offset elimination, straight line subtraction, vector normalization, min/max normalisation and multiplicative scattering correction (Griffith and Anderson 2019). All combinations of these options were iterated to find the best fit, based on the root mean squared error (RMSE). The best performing model was decided on the coefficient of determination and the RMSE for the validation set. This modelling was run across all species and for each separate species.

### 2.5 Modelling FMC through time

Linear mixed-effects modelling was used to understand time and environmental effects on diel leaf FMC. Leaf FMC was used as the response variable and time was always included as a factorial predictor variable to account for leaf FMC being repeatedly measured at eight timepoints in each of the sampling days. Individual tree was included as a random factor in all models. To remove variables that did not contribute to the model, a variable selection process was run, which included sequential removal and testing for collinearity. To judge these differences, Akaike Information Criterion (AIC) and conditional R^2^ values were computed. To find the magnitude and significance of predictor effects, including at different levels of predictors, ANOVA tables of fitted models and the associated coefficients were studied. The pairwise mean estimates were computed, adjusted using the Šidák (1967) method, to compare differences in marginal means within and between predictors. This modelling was done using the software packages lme4, lmerTest, performance, sjPlot, car, multcomp in the R environment (Bates et al. 2015, Kuznetsova et al. 2017, Lüdecke et al. 2021, Lüdecke 2023, Fox & Weisberg 2018, Hothorn et al. 2008, R Core Team 2022).

## 3. Results

### 3.1 Diel leaf FMC

We found leaf FMC had a mean of 121.2 %DW and standard deviation of 21.5 %DW, in a distribution that was slightly positively skewed and included some low outliers (Figure 1). This result shows that over one day and night, FMC of some eucalypt species fluctuated by as much as one fifth, relative to dry weight. We found that the diel range of FMC in individual leaves, across the study, had a mean of 23 %DW, and was as much as 46 %DW or as little as 10 %DW in particular leaves during specific periods (95^th^ and 5^th^ percentile respectively, Figure 2). This range indicates that there is large variation in diel changes amongst species and trees. We don’t report the absolute minima and maxima in the range of diel FMC because a few leaves severely dehydrated to the point of browning in heatwaves.

**Figure 1.**
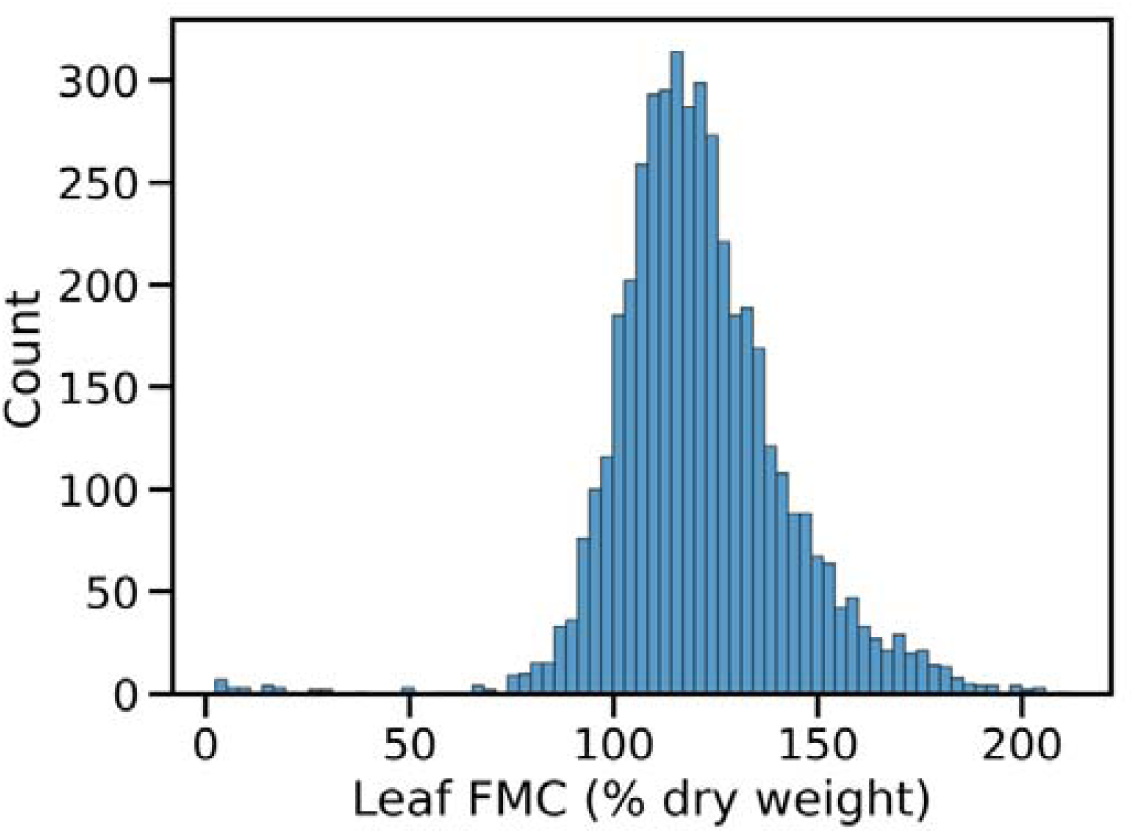
FMC distribution from all leaf measures in all diel periods. FMC predicted using species specific spectral models. Mean = 121.2 % dry weight, std. dev. = 21.5 % dry weight.

**Figure 2.**
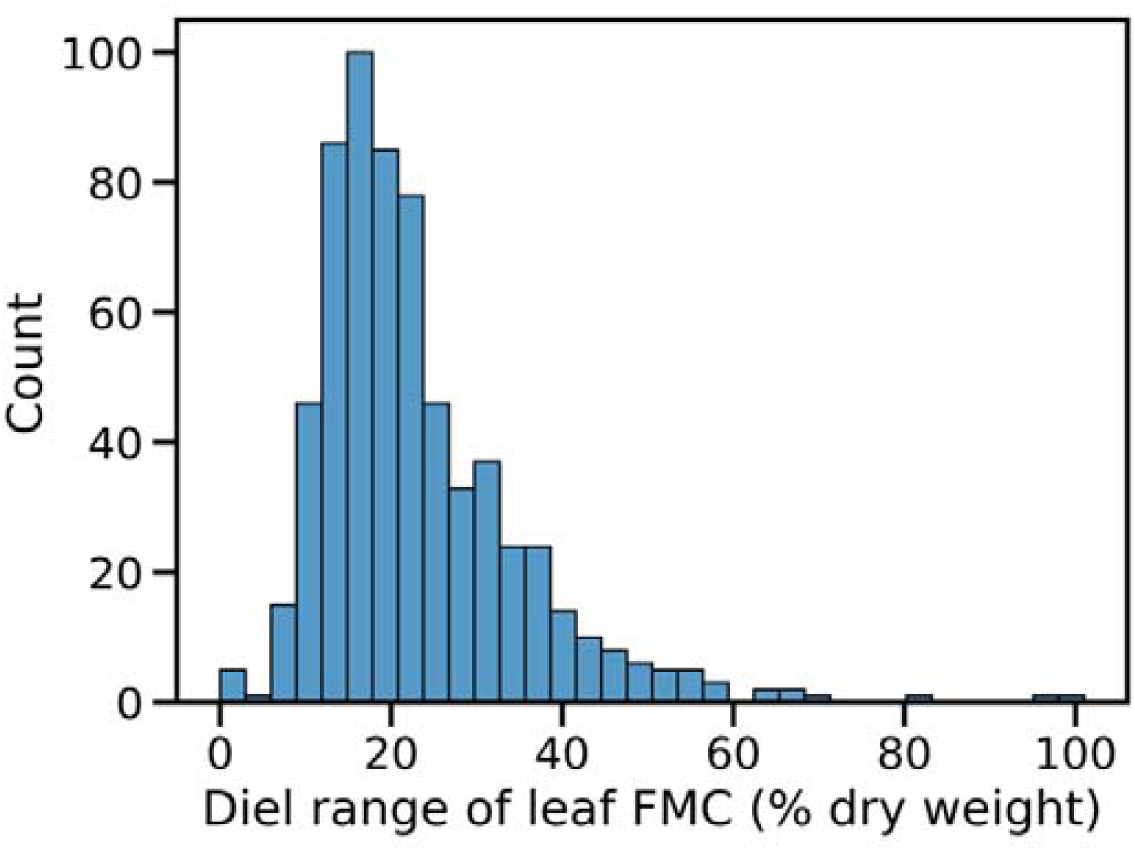
Distribution of the range of FMC of each leaf within each diel period (n=639). Mean = 23.0 % dry weight, median = 20.3 % dry weight.

Amongst species the mean leaf FMC was highest in *E. moluccana* (133.9 %DW), followed by *E. tereticornis* (129.9 %DW), then *E. fibrosa* (116.6 %DW), then *E. crebra* (114.7 %DW) and then *E. sideroxylon* (111.3 %DW). This pattern amongst species was reflected in the range of diel leaf FMC, with largest mean range in *E. moluccana* (32.9 %DW), followed by *E. tereticornis* (21.4 %DW), then *E. fibrosa* (20.8 %DW), then *E. crebra* (20.8 %DW) and then *E. sideroxylon* (19.3 %DW). This shows that some species have ∼40 % greater fluctuation of leaf FMC within the diel period than other species, with higher fluctuation generally corresponding to higher mean FMC. However, *E. tereticornis* had a relatively small diel FMC variation than might be expected based on the species’ relatively high mean FMC.

We found there was a significant effect of diel timepoint on leaf FMC (F-stat.=19.66, *P<*0.0001, Table 3), meaning leaf FMC did change through the day and night. The effect of timepoints on FMC was strongest, negatively, relative to the initial time (07:30), at 13:30 and weakest at 05:00 (marginally insignificant, Table 4). This means that leaf FMC declined through the late morning until 13:30 and then time had a lessening negative effect with the largest reduction in effect (i.e. increase in FMC) between 13:30 and 16:30.

**Table 3.** Analysis of variance table for best fitted model of leaf FMC, without considering all levels of the time factor, which is the 3 hourly diel timepoint (i.e. repeat measures). PAR is photosynthetically active radiation measured by in-house sensors.

|  | FMC |  |  |  |  |  |
| --- | --- | --- | --- | --- | --- | --- |
|  | Sum Sq | Mean Sq | NumDF | DenDF | F value | Pr(>F) |
| Soil Moisture | 67566.96 | 67566.96 | 1.00 | 4053.53 | 454.93 | **<0.001** |
| Temperature | 8334.77 | 8334.77 | 1.00 | 4051.88 | 56.12 | **<0.001** |
| Humidity | 415.97 | 415.97 | 1.00 | 4054.07 | 2.80 | 0.094 |
| PAR Light | 1230.82 | 1230.82 | 1.00 | 4051.16 | 8.29 | 0.004 |
| Time | 20438.28 | 2919.75 | 7.00 | 4050.41 | 19.66 | **<0.001** |
| Species | 7809.56 | 1952.39 | 4.00 | 45.01 | 13.15 | **<0.001** |

**Table 4.**
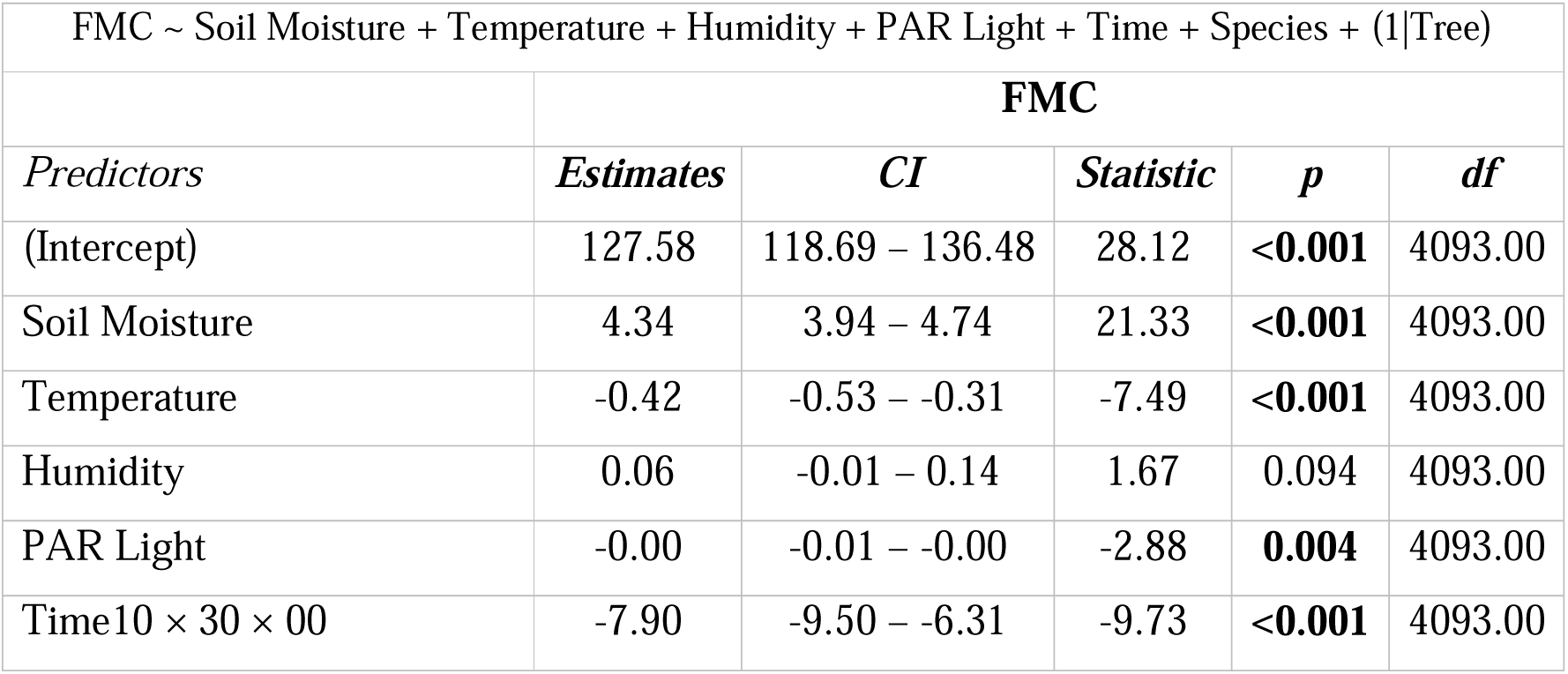
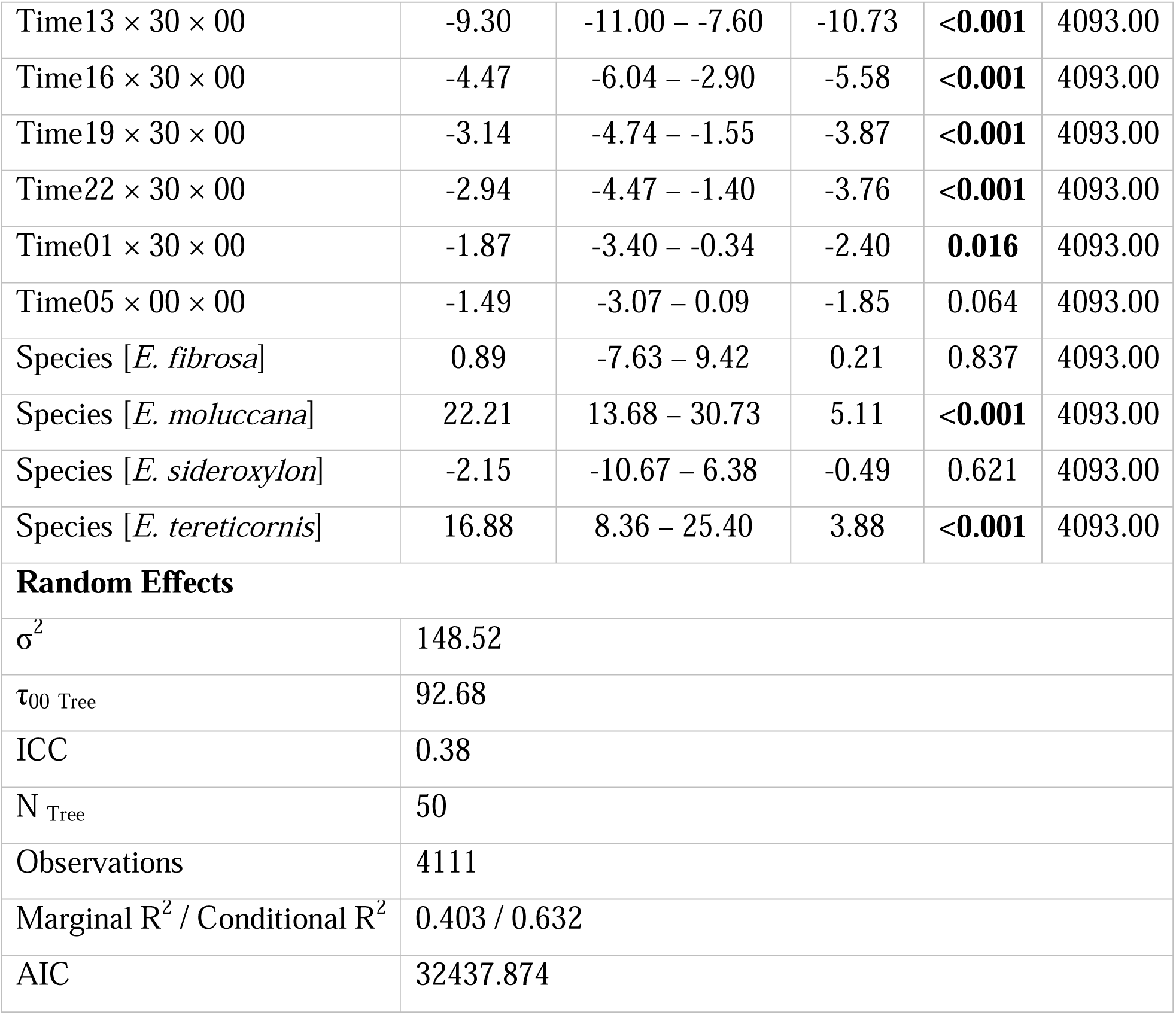
Coefficients with confidence intervals, T-statistics, p-values and degrees of freedom of effects in best fitted model of diel leaf FMC. Estimates for times are relative to 07:30, the first measured each period. Model equation in code format at top of table. Satterthwaite’s degrees of freedom method used for calculating p and df. Note: estimates may appear as zero due to rounding.

| FMC ~ Soil Moisture + Temperature + Humidity + PAR Light + Time + Species + (1\|Tree) |  |  |  |  |  |
| --- | --- | --- | --- | --- | --- |
|  | FMC |  |  |  |  |
| *Predictors* | *Estimates* | *CI* | *Statistic* | *p* | *df* |
| (Intercept) | 127.58 | 118.69 – 136.48 | 28.12 | **<0.001** | 4093.00 |
| Soil Moisture | 4.34 | 3.94 – 4.74 | 21.33 | **<0.001** | 4093.00 |
| Temperature | -0.42 | -0.53 – -0.31 | -7.49 | **<0.001** | 4093.00 |
| Humidity | 0.06 | -0.01 – 0.14 | 1.67 | 0.094 | 4093.00 |
| PAR Light | -0.00 | -0.01 – -0.00 | -2.88 | **0.004** | 4093.00 |
| Time10 × 30 × 00 | -7.90 | -9.50 – -6.31 | -9.73 | **<0.001** | 4093.00 |
| Time13 × 30 × 00 | -9.30 | -11.00 – -7.60 | -10.73 | <0.001 | 4093.00 |
| Time16 × 30 × 00 | -4.47 | -6.04 – -2.90 | -5.58 | <0.001 | 4093.00 |
| Time19 × 30 × 00 | -3.14 | -4.74 – -1.55 | -3.87 | <0.001 | 4093.00 |
| Time22 × 30 × 00 | -2.94 | -4.47 – -1.40 | -3.76 | <0.001 | 4093.00 |
| Time01 × 30 × 00 | -1.87 | -3.40 – -0.34 | -2.40 | 0.016 | 4093.00 |
| Time05 × 00 × 00 | -1.49 | -3.07 – 0.09 | -1.85 | 0.064 | 4093.00 |
| Species [*E. fibrosa*] | 0.89 | -7.63 – 9.42 | 0.21 | 0.837 | 4093.00 |
| Species [*E. moluccana*] | 22.21 | 13.68 – 30.73 | 5.11 | <0.001 | 4093.00 |
| Species [*E. sideroxylon*] | -2.15 | -10.67 – 6.38 | -0.49 | 0.621 | 4093.00 |
| Species [*E. tereticornis*] | 16.88 | 8.36 – 25.40 | 3.88 | <0.001 | 4093.00 |
| **Random Effects** |  |  |  |  |  |
| $\sigma^2$ | 148.52 | | | | |
| $\tau_{00 \text{ Tree}}$ | 92.68 | | | | |
| ICC | 0.38 |  |  |  |  |
| N<sub>Tree</sub> | 50 |  |  |  |  |
| Observations | 4111 |  |  |  |  |
| Marginal R<sup>2</sup> / Conditional R<sup>2</sup> | 0.403 / 0.632 |  |  |  |  |
| AIC | 32437.874 |  |  |  |  |

The lowest marginal mean of FMC occurred at 13:30, confirming our first hypothesis (H_1_), that minimum FMC occurs during the early afternoon, however this was not different to the marginal FMC mean earlier at 10:30. This indicates that FMC changes the fastest between 7:30 and 10:30 at which point it is at the lowest levels through 13:30. There was no difference in FMC marginal means between the times of 7:30, and the following 1:30 and 5:00 (Figure 3). This shows that FMC recovered to previous morning (7:30) levels by 1:30 at night, on average. However, FMC reaches a level statistically equivalent to this recovery time by 19:30, although the mean FMC value did continue to rise, up to the recovery time. These results contradicted our second hypothesis (H_2_) that leaf FMC would recover later in the early morning, by 5:00.

**Figure 3.**
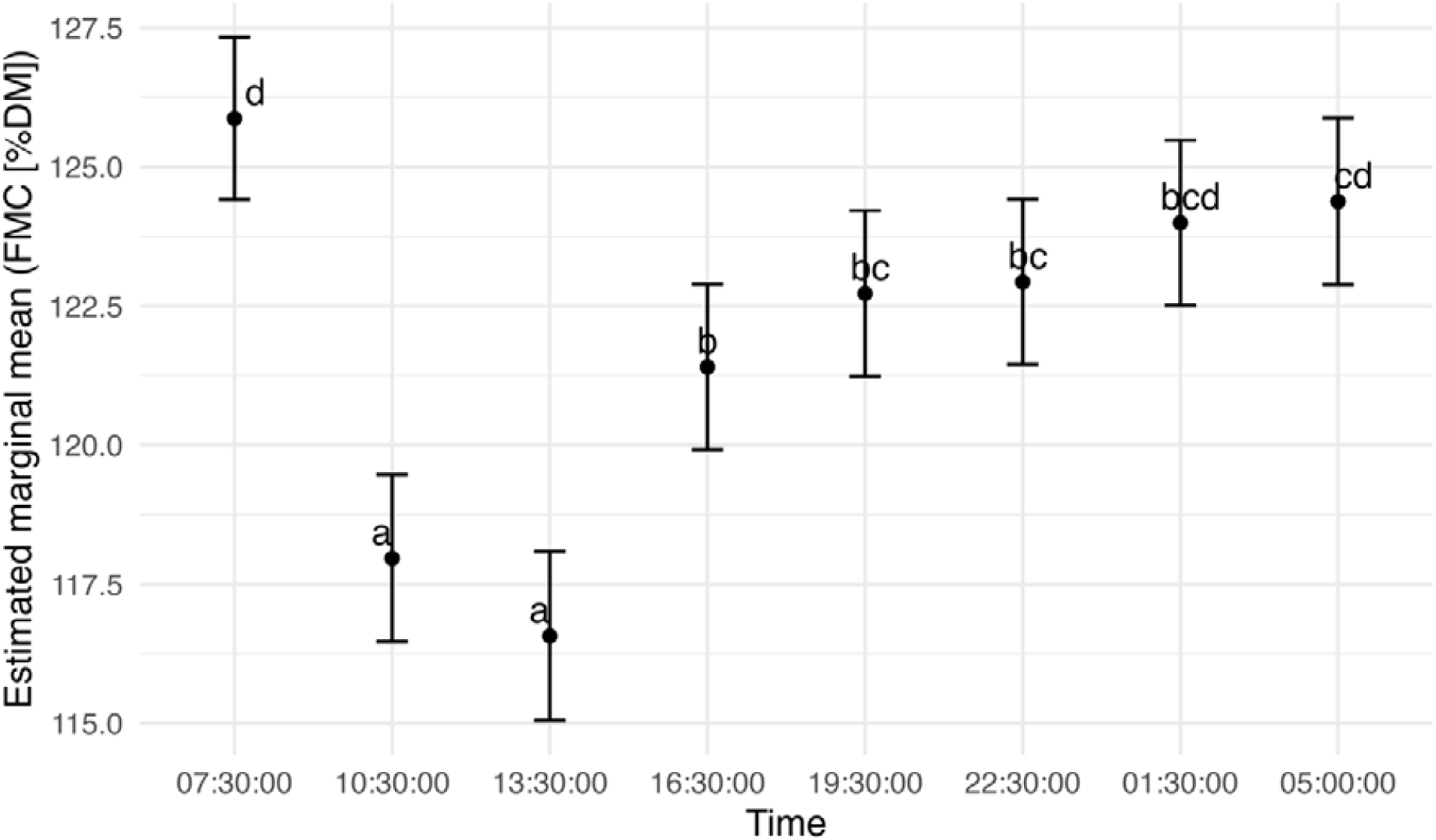
Estimated marginal means and standard errors of FMC across time, from best fitted model, including times that are not significantly different, as indicated by shared letters.

We found there was a significant effect on leaf FMC of tested environmental variables (all *P<*0.001), except elevated CO_2_ which had a slightly negative but insignificant effect on FMC and was therefore excluded from models. Humidity had a weakly significant effect though it contributed to a better model with lower AIC, than not including it (Table 3). These results show there was a change in leaf FMC due to the changes in glasshouse conditions (Supplementary Table S6). We also found that a model including VPD did not perform as well as the model including instead, temperature and humidity, from which VPD is derived (R^2^= 0.622 vs 0.632, AIC= 32437.874 vs 32543.151, respectively). These results shows that changes to atmospheric moisture had a smaller effect on leaf FMC relative to other glasshouse variables.

We found a significant interaction of time and drought on leaf FMC (F-stat. = 2.85, *P=*0.005, 3 of 6 diel measurement periods were droughted). In drought there is no difference between FMC at 7:30 and 19:30 and without drought FMC is no different to 7:30 by 1:30 (Figure 4). The largest average difference between timepoints in FMC across time was 12 % in drought and 7 % without drought. These results indicate that during drought there is a smaller decline in FMC during the day, which is recovered more quickly at night, compared to non-drought conditions, thus contradicting our third hypothesis.

**Figure 4.**
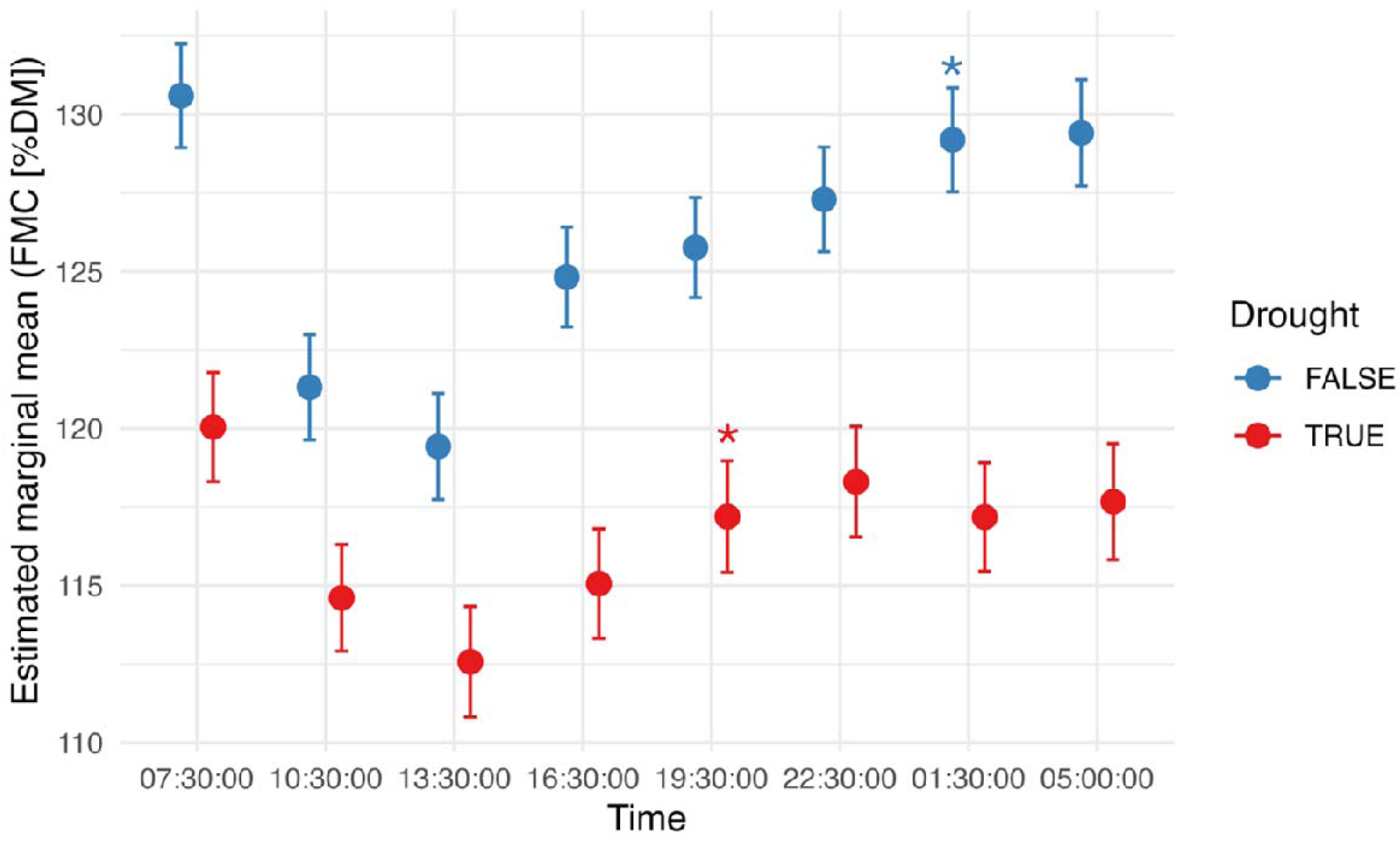
Estimated marginal means and standard errors of FMC across time, from best fitted model, in drought (red) or non-drought (blue) conditions. Asterisks indicate the time at which there is no difference in FMC to that of the initial time, 07:30. Data points are shown to be shifted before or after each timepoint but correspond to the nearest timepoint indicated.

We found that FMC did not differ among *E. crebra*, *E. fibrosa*, *E. sideroxylon* but was significantly less (*P*<0.01) than in *E. tereticornis* and *E. moluccana*, which were no different to each other. *E. moluccana* had the highest mean and *E. crebra* the lowest. These results indicate that leaf FMC is variable amongst *Eucalyptus* species, including that FMC differs amongst *Eucalyptus* species.

There was a significant interaction between time and species (F-stat. = 3.42, *P<*0.0001). By 16:30, leaf FMC levels were no different to the initial, morning time (7:30) in *E. fibrosa*, *E. moluccana* and *E. sideroxylon*. For *E. crebra* in the following timepoint (19:30) there was no difference to the initial time, and for *E. tereticornis* this recovery occurred later again at 1:30 for (Figure 5).

**Figure 5.**
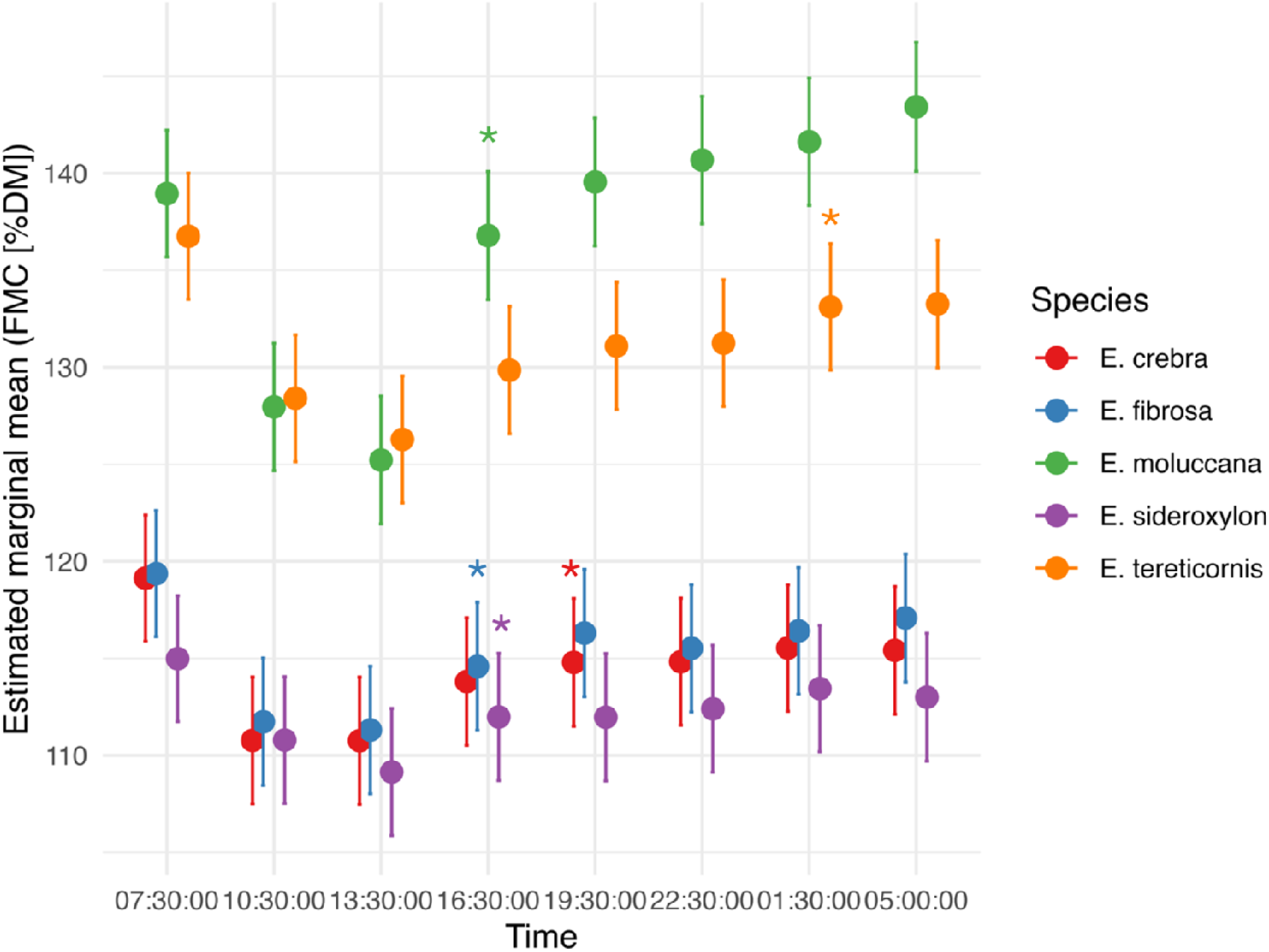
Estimated marginal means and standard errors of foliar moisture content (FMC) across time, from best fitted model, per species. Asterisks indicate the time at which there is no difference in FMC to that of the initial time, 07:30. Input data is from all six diel periods, including heatwaves, drought and well-watered conditions. Data points are shown to be shifted before or after each timepoint but correspond to the nearest timepoint indicated.

### 3.2 Calibration of NIRS FMC model

We found that the calibration dataset from bench drying leaves had a mean FMC of 81.51 %DW, with a minimum of 8.6 and maximum of 233.5 %DW (Figure 6). This result shows that the bench drying reduced leaf moisture, and that there was a positive skew in the distribution of FMC in the calibration dataset.

**Figure 6.**
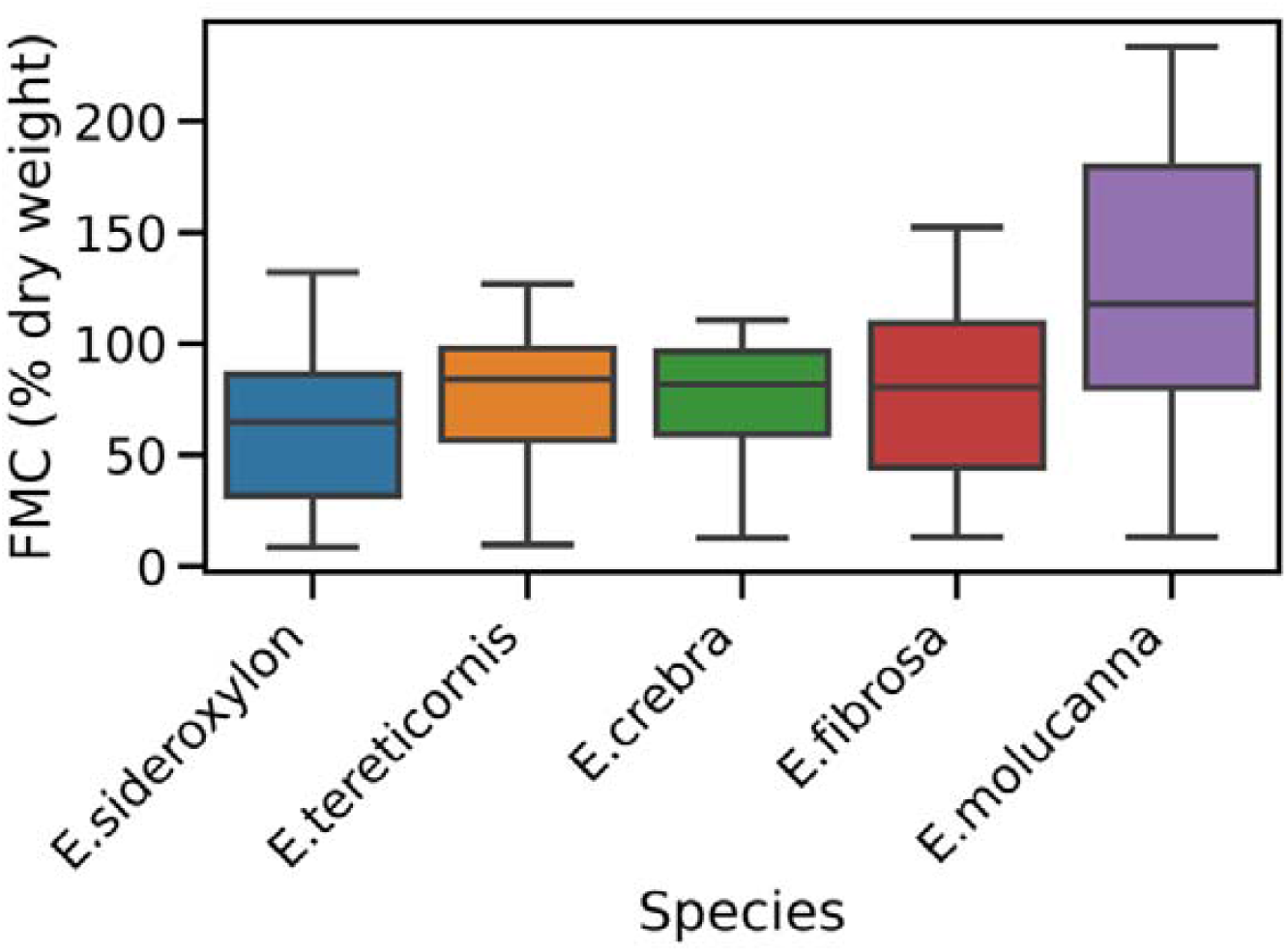
Quartiles and extremes of leaf FMC of repeat measures during bench drying by species. Species sorted by respective means.

We found that the best NIRS model of FMC had high predictive power (r^2^_cal_>0.9) and errors of 10 %DW or less (Figure 7, Figure 8, Table 5). This means that reflectance data from a leaf predicts the moisture content of that leaf with high accuracy. The species-specific models had lower predictive power than the cross-species model (data not shown). The second validation produced a less accurate result (r^2^=0.74) with slightly higher errors (RMSE = 12.46 %DW, Figure 9). This result suggests that there may be a lack of representation in the calibration dataset. The best NIRS model used only near infrared wavelengths between 1600 to 2400 nm and used min/max normalization of leaf averaged spectra (Supplementary Table S7). This result suggests that the water content in leaves affects light in the longer part of the short-wave infrared portion of the spectrum.

**Figure 7.**
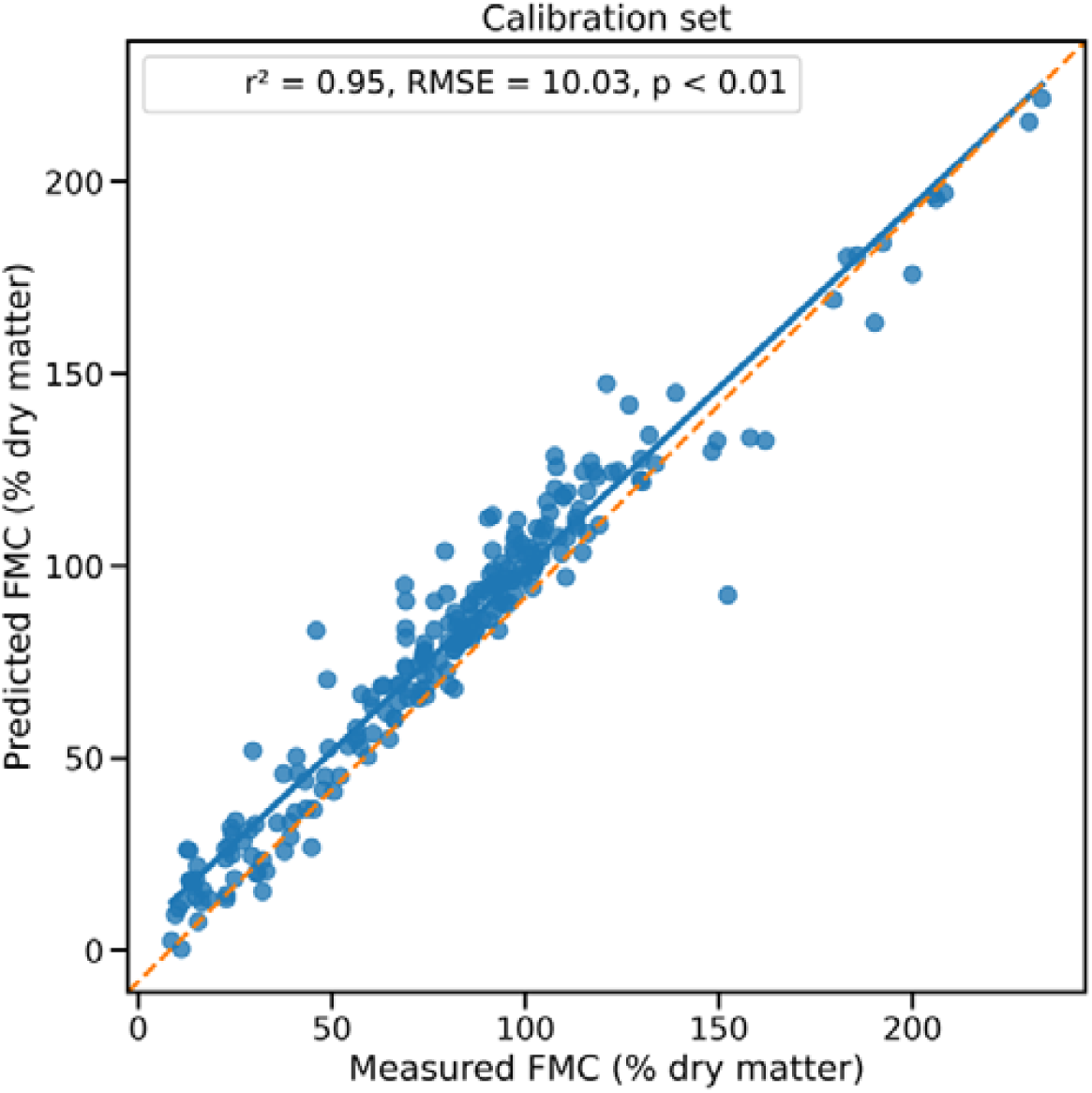
Prediction of FMC on the calibration set (n=228) from the fitted, best model. Prediction plotted against FMC measure taken during the dry down process.

**Figure 8.**
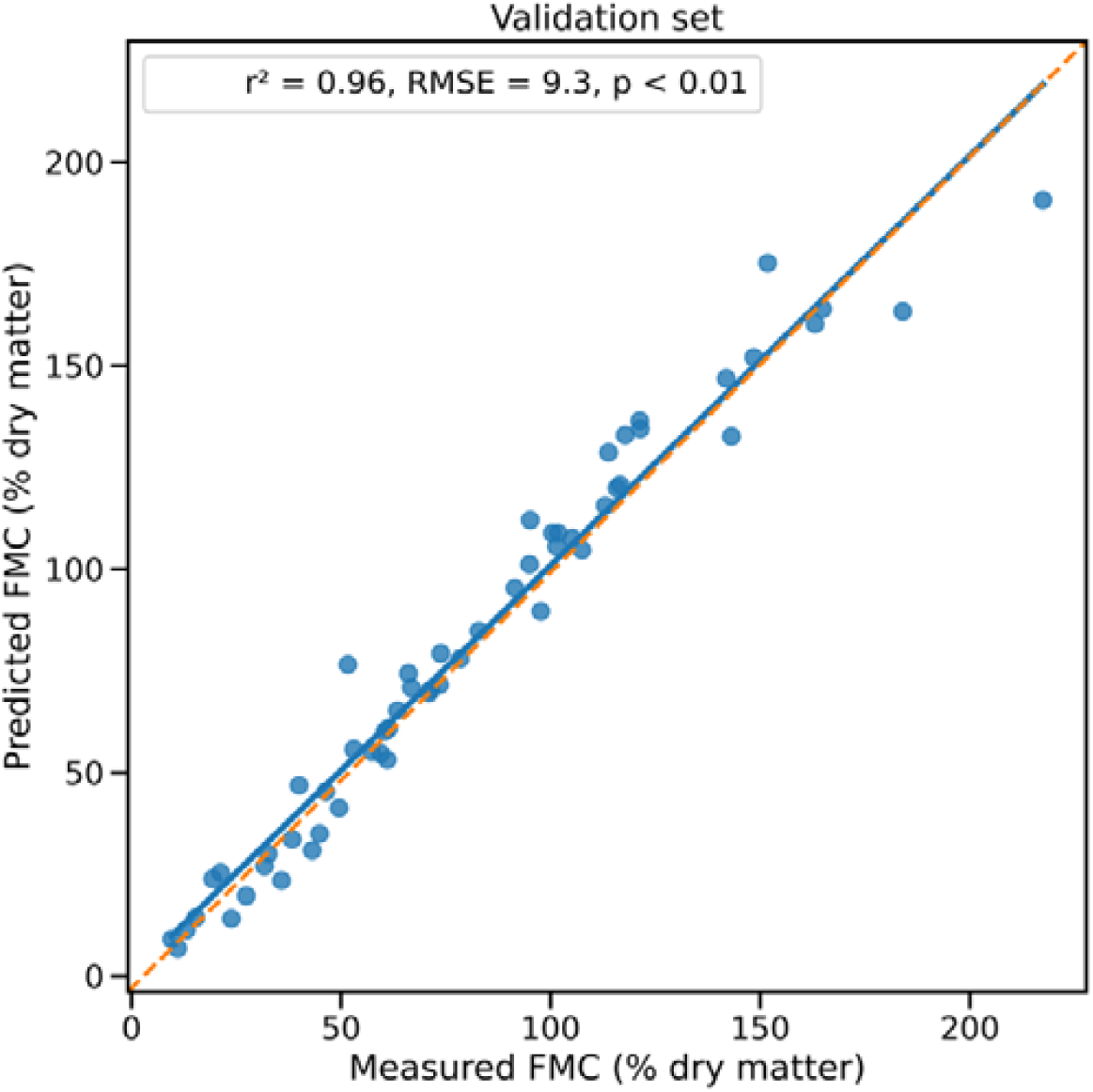
Prediction of FMC on the validation set (n=57) from the fitted, best model. Prediction plotted against FMC measure taken during the dry down process.

**Figure 9.**
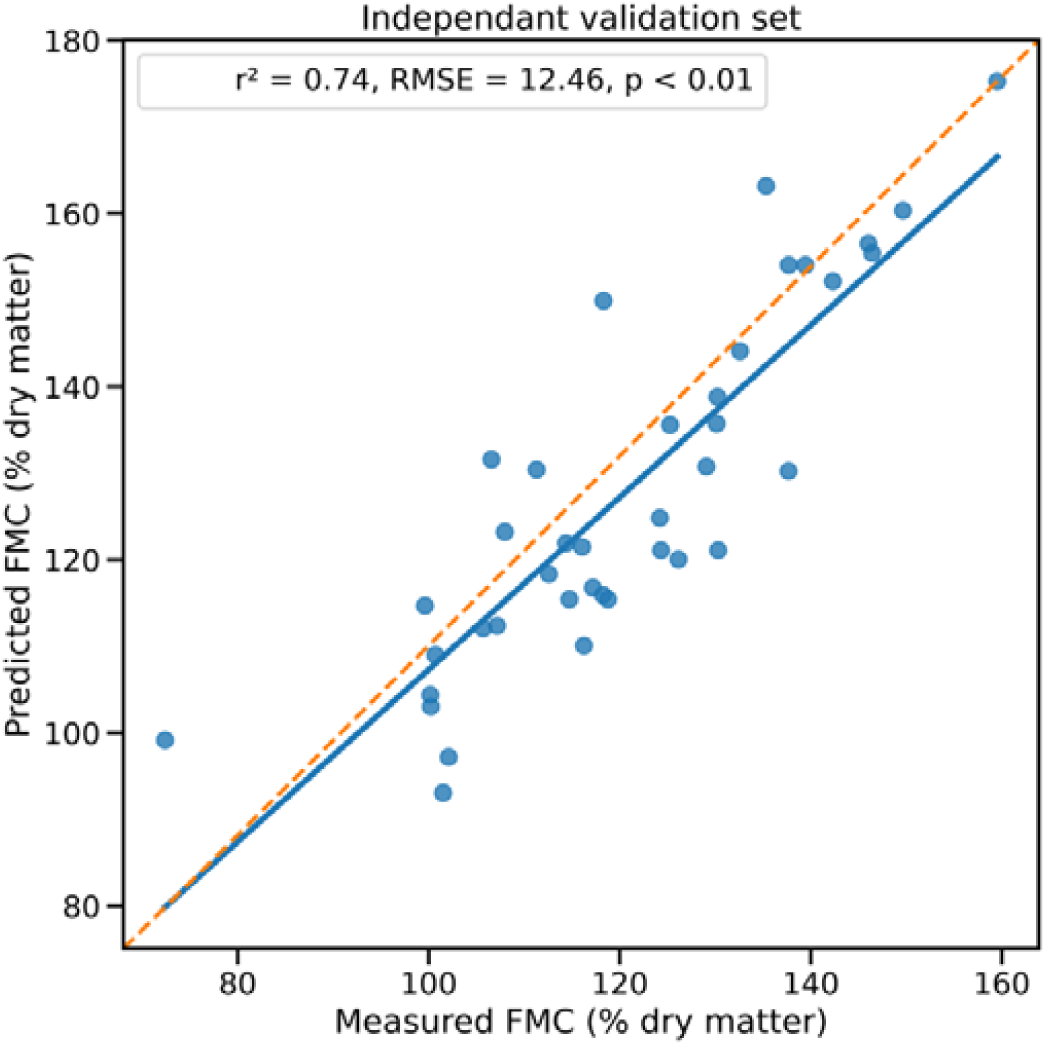
Prediction of leaf FMC on second validation set (n=39) from the fitted, best model. Prediction plotted against FMC measure taken during the glasshouse experiment.

**Table 5.** Errors and accuracy of the predictive spectral model of FMC from leaf bench drying samples. n represent the average of 3 spectra per leaf.

| Species | n | Calibration |  |  |  | Validation |  |  |
| --- | --- | --- | --- | --- | --- | --- | --- | --- |
|  |  | n | Factors | RMSE<sub>p</sub> | r<sup>2</sup> | n | RMSE<sub>p</sub> | r<sup>2</sup> |
| All (*E. crebra*, *E. moluccana*, *E. fibrosa*, *E. tereticornis*, *E. sideroxylon*) | 285 | 228 | 8 | 10.03 | 0.95 | 57 | 9.29 | 0.96 |

## 4. Discussion

In this study we found diel ranges in leaf FMC of 23 %DW on average across species and individuals, which is substantial in the context of koala forage. We focus the discussion on our results of diel FMC range of individual leaves rather than the mean, modelled range across leaves because koalas experience the absolute value of moisture in individual leaves not a statistical average. Other studies have found that seasonal differences in mean leaf FMC of koala habitats are 5, 12 or 18 %DW (Hume & Esson 1993, Melzer 1994, Ellis et al. 1995). These seasonal changes were associated with a preference for different species across the season (Ellis et al. 1995), and for different trees of varying leaf moisture levels of the same species (Melzer 1994). The magnitudes of these seasonal changes are less than the changes we found within the course of an average day and night. Therefore, we suggest that diel variation in FMC could be a factor in forage selection by koalas during high temperatures when water demand for evaporative cooling is high. This will be the case especially for koalas in habitats lacking open water which could be used to supplement dietary water (Mella et al. 2019). Furthermore, the diel variation in FMC is additional to seasonal or longer scale variation.

We observed an effect of time on leaf FMC that was different across tree species, which may allow koala foraging in a similar way to that which (Ellis et al. 1995) observed between seasons, with water intake being increased by choosing different species through time. Therefore, koala preference may be for trees which maintain a higher minimum diel FMC, such as *E. tereticornis* rather than *E. moluccana* which declined further despite having a higher morning FMC level, to increase the chance across the day of gaining sufficient water from the diet. Similar preferences were observed by Wu et al. (2012) where the lower FMC, ironbark *Eucalyptus melanophloia* became an important habitat tree during drought compared to tree species which rely more greatly on soil moisture along creeks or rivers which dried up (e.g. *Eucalyptus coolabah* and *Eucalyptus camaldulensis*).

Differences amongst species in diel FMC generally reflected the species’ climate of origin (Belbin et al. 2021). Species with a range covering more arid climates had on average, the lowest means and diel ranges of FMC while species from mainly mesic climates had the highest means with a larger diel range (*E. sideroxylon* and *E. tereticornis*, respectively). Species covering arid climates regulate their water use and loss more strictly than species experiencing high water availability (Coleman et al. 2023), including through increased stomatal sensitivity to VPD (Bourne, Haigh and Ellsworth 2015). However, the pattern observed in mean and range of diel FMC was not directly reflected in the diel trend, with the two most mesic species recovering to morning levels at different times (*E. moluccana* at 16:30, E*. tereticornis* at 1:30). This result may have been affected by the different sizes of the trees, *E. moluccana* were larger and older than other species with greater leaf area that may have increased the rate of water uptake from the soil to leaves. Differences in diel FMC amongst species is important because of the preference koalas have for a small set of species, in most populations, due largely to nutritional attributes and plant secondary metabolites in leaves which are foraging deterrents (Melzer et al. 2014, Moore et al. 2010). Au (2018) found that *E. tereticornis* contained the highest level of formylated phloroglucinol compounds (FPCs) amongst our study species, the primary deterrent in the shared sub-genera (*Symphyomyrtus*). This and our finding of slower recovery time suggests *E. tereticornis* may be poorer quality in terms of toxicity and dryness during evening foraging in heatwaves when water demand is high, compared to our other tree species, assuming a similar species composition to that used here (Moore et al. 2010). However, *E. tereticornis* is known to be a preferred species, containing low concentrations of FPCs in some habitats (Youngentob, Marsh and Skewes 2021). The potential impact of diel FMC on habitat quality also depends on several other factors. These include competition amongst koalas (Marsh et al. 2013), microclimate variation within habitats (Ellis et al. 2010) and other forage selection pressures like the composition of all nutrients (Beale et al. 2018). Despite these potential interactions the diel FMC range and variation amongst species we observed was substantial and may have some influence on foraging choices.

While the diel FMC magnitude is substantial, the FMC minima does not generally occur during usual koala foraging times. There is evidence that koalas forage preferentially in the evening to lessen heat load in warm environments (Adam et al. 2021). Our results suggest that this preference also avoids the minimum in diel FMC (i.e. ∼13:30), so the degree of impact of FMC on behaviour or physiology may be reduced. However, Adam et al. (2020) found that koala diel body temperature on the hottest day in their study was above long-term average until 4:00 or 5:00 the following morning, and in one case it never approached average, increasing the need for water to increase evaporative cooling. In such extremes, the recovery of FMC after midnight (1:30) to previous morning levels, as found here, may be important for koala thermoregulation, in that foraging may be adjusted to this time or later, to gain more water from moister leaves. We are assuming here that free water is not available, as is the case in many habitats during extreme heat (Mella et al. 2019). Another factor in feeding time may the thermal cost of feeding (Beale et al. 2018), which presumably lessens, relatively, as the night becomes colder. Furthermore, the importance of night-time recovery of FMC may increase if the FMC minima is delayed during hot and dry conditions. We are unable to answer that question in this study due to the relatively few sampling days of contrasting glasshouses conditions. Therefore, discerning the relative, temporal effects of both FMC and temperature on koala feeding times is a promising direction for new research.

Although koala foraging tends to occur other than during the diel FMC minima observed here, the rate of return to FMC maxima may still be important in certain conditions if koalas can forage for moisture. There is evidence in the koala genome for the ability to sense moisture in forage, amongst other nutritive and toxic components (Johnson et al. 2018). The authors of that study suggested koalas may be able to sense water because of the aquaporin 5 gene being functionally duplicated. This genetic feature may allow behavioural responses to FMC changes in habitat tree leaves, such as the changes through the average diel range of 23 %DW found in this study. It also recognised that there is an innate thirst drive in other animal species (Moore & Foley 2000). Such behavioural responses to forage moisture have been recorded in koalas between seasons. In a buffet feeding experiment (Melzer 1994) found there are thresholds in forage moisture levels below which koalas choose different trees to substantially reduce intake of that forage, and that these thresholds differed by 12 %DW between seasons. The seasonal differences were an increased mean maximum air temperature and mean VPD (21 % and 33 %, respectively). Given the behavioural and genetic evidence for moisture selection of forage, our results reveal a magnitude of diel change that may drive the need for such selection during hot and dry conditions, even over daily timescales.

Phillips et al. (2010) observed diel stomatal conductance of two of our study species to peak between 10:00 and 13:00, the same times over which we observed FMC to be lowest in these species (their Figure 3, *E. tereticornis*, *E. sideroxylon*). This indicates that minimum FMC levels are limited by the rates of water loss from leaves. The return of stomatal conductance to morning levels before sunset is similar but somewhat faster than the recovery we observed for FMC, though there FMC did start to increase before sunset. The difference between stomatal conductance and FMC recovery was least similar in *E. tereticornis* which on average recovered FMC later than other species, at 01:30. This difference indicates that leaf refill with water can take some time after stomata close.

The large average diel variation and leaf diel range in FMC is potentially a source of error in instantaneous remote sensing estimates of FMC that are generally taken to be a daily value, often due to infrequent satellite revisits (e.g. Kotzur et al. 2025). Satellite measures, taken outside the 10:30 to 16:30 window, are likely to overpredict minimum FMC. Minimum FMC is often of interest, both in animal ecology where extreme low values may limit species’ thermo-physiology and distribution (Briscoe et al. 2016) and in fire ecology where low thresholds are related to fire behaviour (Nolan et al. 2016). However, this period is when many optical satellite sensors are taking measurements to utilise the high sun position (i.e. a sun-synchronous orbit), limiting the difference with diel FMC minima as observed here. The diel change in FMC may also be important for measures taken in the field including remote sensors or destructive measures. Measures of FMC taken in the morning must be done close to dawn if the maximum is of interest, because the decline in FMC early in the day is rapid compared to the later increase. Existing methods in fire ecology and plant physiology tend to account for this by sampling prior to dawn for the maximum value of water balance variables (Nolan et al. 2018). Our results suggest that sampling could alternatively be done late at night to achieve a similar value, however this average result cannot be guaranteed across all species and conditions. This finding that FMC recovers before the predawn measurement, disproves our second hypothesis and shows that trees are ready to resume leaf hydration quickly, on average, including when experiencing a mix of neutral and demanding atmospheres and varying soil hydration. Future work should test if there is a greater time lag, in this respect, for larger trees which have a greater volume of water vessels (Choat et al. 2018).

We were able to predict live leaf FMC with high accuracy, which has rarely been done with NIRS despite many studies of water content (i.e. % fresh weight) and not at this temporal resolution. The accuracy achieved here confirms that the dry weight which is the denominator in the FMC calculation is accounted for in the spectral information captured in PLS models from full range spectra (Ceccato 2001). The model performed better than previous NIRS models of leaf water potential and relative water content of *Eucalyptus* (Yang et al. 2017), likely because those measures are functional and relative, respectively, whereas FMC is an absolute measure of leaf water which drives the strong spectral response of the water absorbance features (Datt 1999). Our second validation set did provide slightly larger errors and lower accuracy than the initial validation set. This shows the performance of NIRS models may be variable even when predictions are made on the same leaves as used in model calibration, something of note also if looking to make predictions using reflectance from another remote sensor, field or satellite based, which introduces further, spectral variability as well as reducing the range and resolution available for prediction. This degradation in performance may be explained by the extreme conditions imposed on leaves of the second validation set, throughout the experiment, which damaged the hydraulic vessels in the trees (Choat et al. 2018) to a degree that caused some leaves to completely desiccate during the experiment. Though we avoided choosing desiccated leaves, for the second calibration set, leaves may still have been internally damaged. The poorer performance with the second validation set could also partly be explained by the negative skew in the calibration set (Figure 6) relative to the second validation set. When drying leaves to make the calibration set, FMC declined rapidly at the beginning, increasing the labour needed for collecting moist samples. Whereas the second validation set was comprised of, potentially damaged though, relatively moist leaves measured immediately after removal from the tree. The decreased accuracy of the second validation could be improved by increasing the sampling effort at the beginning of the dry down process, and by enlarging the calibration set overall to include greater temporal and leaf structure variation. For analysing diel FMC of live leaves in the glasshouse the NIRS model was accurate, and this result provides a great opportunity via non-destructive, repeat leaf measures for future studies of FMC and plant water relations. Such studies could be based on airborne hyperspectral reflectance data, with a new validation of the FMC model considering potentially large effects of atmospheric interference, spectral resolution and range, and non-target signals, to quantify tree scale water availability to koalas and could reveal composition of forage quality within habitats in terms of heat refugia. The quantification of fine spatial scale and daily water availability has been undertaken in previous work (Kotzur et al. 2025), revealing high variability in extreme drought response within landscapes (*Kotzur et al. unpublished*). Application of the finer temporal trend identified in this study to such satellite-based models of FMC, which typically use satellites of daily revisit times, will further refine the prediction of potential heat-drought refugia in forest environments by revealing sub-daily FMC trends. The sub-daily trends could be put in terms of atmospheric demand and soil moisture to spatially apply variation on to the daily FMC measures via existing climatic and landscape variables. A similar result of sub-daily fuel moisture estimates has recently been achieved using a different approach in fire ecology, utilising geostationary satellite to produce 10 min predictions (Quan et al. 2024). Such work highlights the potential application of integrating the sub-daily and daily FMC trends. The resulting data could be combined with koala tracking to analyse forage selection and water concentrations at different air temperatures to understand habitat selection through extremes.

The study design carries several other implications for our results. Firstly, the NIRS model of leaf FMC developed here is applicable to mature trees in the field using a similar spectrometer, though there may be differences in the magnitude of diel variation. This may be driven by differences in the hydraulic structure between small, young and large, old trees. Ageing in trees tends to increase the average leaf mass per area (LMA, Thomas & Winner 2002), including in *Eucalyptus* (Wujeska-Klause et al. 2019). FMC in mature *Eucalyptus* woodland has been shown to decrease with increasing specific leaf area (SLA, the inverse of LMA) (Griebel et al. 2023). Therefore, we could expect our estimates of diel leaf FMC in relatively young potted saplings to overestimate that of mature trees due to relative increases in dry weight with tree age, and because some leaves measured may have been in a juvenile life stage. However, the leaves studied here were all fully expanded leaves with leaf ages up to 12 months, so potential overestimates may be reduced for similarly mature leaves of older trees in the field. Secondly, the leaf FMC response we observed may have been influenced by the potted and watered study design. We watered plants twice per day at 09:00 and 21:00 using an automated dripper system. The trees were likely to take up available water quickly because of their young age, small internal water storage and the limited volume of potted soil. This response may have temporarily increased the FMC signal, after watering. However, we selected watering times immediately after measurements to allow any pulse in leaf FMC to dissipate. Furthermore, we did not see a response to watering in the diel trend, although there was no control for this so we cannot be certain. Future studies of diel FMC should consider using a temporally constant water supply such as capillary mats (Semananda et al. 2018), to prevent any water uptake effect. Despite these concerns the design allowed high temporal resolution measurement of diel patterns of leaf FMC in some koala preferred *Eucalyptus sp.* trees.

## Conclusion

This study revealed that diel variation in FMC can be substantial through 24 h and is larger than important FMC variation in koala forage from published data, across different scales. It was also shown that diel FMC usually recovered 1.5 h after midnight after a low at 13:30 in the afternoon. However, there was variation amongst species, which may be consequential for koala water intake during evening foraging in high temperatures. Such information is important for understanding koala thermo-physiology in the field and for the conservation of koalas in environments where heat induced water stress is threatening survival. This study also demonstrated a spectroscopy method applicable to monitoring water quality of forage selected by herbivores in the field or in captive studies.

