## Supplementary for "Diel foliar moisture content recovery time occurs soon after midnight in *Eucalyptus species* of koala (*Phascolarctos cinereus*) habitats"

Glasshouse conditions

Table S6. Temperature and humidity targets and actual mean values, including calculated VPD, during 7 different diel sampling periods.

| Chamber | Date_start | Date_end | Humidity Target(%Rh)_mean | Humidity(%Rh)_mean | vpd(kPa)_mean | Temperature Target(oC) | Temperature(oC) |
| --- | --- | --- | --- | --- | --- | --- | --- |
| G09 | 2022-07-28 06:30:00+10:00 | 2022-07-29 05:30:00+10:00 | 60 | 58.6 | 1.08131769 | 20.6167509 | 20.81494585 |
| G09 | 2022-09-05 06:30:00+10:00 | 2022-09-06 05:30:00+10:00 | 50 | 41.1 | 3.092064982 | 33.42187726 | 32.78050542 |
| G09 | 2022-09-06 06:30:00+10:00 | 2022-09-07 05:30:00+10:00 | 30.50577617 | 39.9 | 3.219173285 | 33.42187726 | 31.86310469 |
| G09 | 2022-09-13 06:30:00+10:00 | 2022-09-14 05:30:00+10:00 | 60 | 59.0 | 1.068090253 | 20.6167509 | 20.88761733 |
| G09 | 2022-11-02 07:30:00+11:00 | 2022-11-03 06:30:00+11:00 | 60 | 54.7 | 1.179061372 | 20.55097473 | 20.81617329 |
| G09 | 2022-11-06 07:30:00+11:00 | 2022-11-07 06:30:00+11:00 | 50 | 46.4 | 2.930729242 | 33.24859206 | 33.00826715 |
| G09 | 2022-11-08 07:30:00+11:00 | 2022-11-09 06:30:00+11:00 | 30.50577617 | 46.16895307 | 2.84000361 | 32.8067148 | 31.0767148 |
| G10 | 2022-07-28 06:30:00+10:00 | 2022-07-29 05:30:00+10:00 | 60 | 57.2 | 1.116379061 | 20.6167509 | 20.81761733 |
| G10 | 2022-09-05 06:30:00+10:00 | 2022-09-06 05:30:00+10:00 | 50 | 41.06714801 | 3.123184116 | 33.42187726 | 32.95039711 |
| G10 | 2022-09-06 06:30:00+10:00 | 2022-09-07 05:30:00+10:00 | 30.50577617 | 38.69422383 | 3.290151625 | 33.42187726 | 32.16093863 |
| G10 | 2022-09-13 06:30:00+10:00 | 2022-09-14 05:30:00+10:00 | 60 | 58.00866426 | 1.093942238 | 20.6167509 | 20.91971119 |
| G10 | 2022-11-02 07:30:00+11:00 | 2022-11-03 06:30:00+11:00 | 60 | 53.35054152 | 1.211465704 | 20.55097473 | 20.85126354 |
| G10 | 2022-11-06 07:30:00+11:00 | 2022-11-07 06:30:00+11:00 | 50 | 45.55776173 | 2.95866787 | 33.24859206 | 33.01584838 |
| G10 | 2022-11-08 07:30:00+11:00 | 2022-11-09 06:30:00+11:00 | 30.50577617 | 43.05451264 | 3.055805054 | 32.8067148 | 31.65740072 |

Calibration of NIRS model

Table S7. Results from the optimisation of the partial least square models of spectra and FMC. RMSEP: root mean square error of prediction, Rank: the number of latent variables in the PLS regression, Regions: spectral regions considered in the model, Preprocessing: spectral transformation used with the option of first and second derivatives (D1F and D2F), constant offset elimination (COE), straight line subtraction (SLS), vector normalization (SNV), min/max normalization (MMN), multiplicative scattering correction (MSC).

| **Model no.** | **RMSEP** | **No. of terms** | **Regions (nm)** | **Pre-processing** |
| --- | --- | --- | --- | --- |
| **1** | 7.87 | 9 | 1600 - 2000, 2000 - 2400 | MMN |
| **2** | 8.72 | 9 | 400 - 600, 2000 - 2400 | SNV |
| **3** | 8.82 | 9 | 400 - 600, 1600 - 2000, 2000 - 2400 | SNV |
| **4** | 9.01 | 10 | 1600 - 2000 | MMN |
| **5** | 9.42 | 9 | 400 - 600, 600 - 800, 1200 - 1600, 1600 - 2000, 2000 - 2400 | SNV |
